# scLANTERN: High-Throughput Retrospective Lineage Tracing via Full-Length Single-Cell Transcriptomics and Expressed Repeat Variation

**DOI:** 10.64898/2026.08.25.747112

**Authors:** Liming Tao, Jack Kamm, Yuntian Fu, Duc Nguyen, Nicolò Riggi

**Affiliations:** Department of Cell and Tissue Genomics, Genentech Inc, South San Francisco, CA, USA; Computational Sciences Center of Excellence, Genentech Inc, South San Francisco, CA, USA; Department of Discovery Oncology, Genentech, Inc, South San Francisco, CA, USA; Department of AI Biology & Translation, Genentech, Inc, South San Francisco, CA, USA

## Abstract

Understanding the lineage relationships among individual cells is a key pursuit of modern biology, essential for unraveling the complexities of developmental processes and the adaptive mechanisms of disease progression, particularly in oncology. Retrospective single-cell clonal tracing has emerged as a transformative approach, offering a unique window into the evolutionary trajectories of cancer within clinical samples. While short-read single-cell transcriptomics (scRNA-seq) has revolutionized our ability to map cell states across human tumor atlases, it remains fundamentally limited in its capacity to link these states with high-resolution genomic alterations and the evolutionary trajectories inferred from these natural variants. Integrating somatic mutation discovery with transcriptomic profiles at single-cell resolution often requires separate, costly, and low-throughput genomic assays. Furthermore, existing methods frequently rely on exogenous genetic labeling or are restricted to short-read sequencing, which typically fails to resolve complex genomic rearrangements, large indels, or variations within highly repetitive regions—such as short tandem repeats (STRs)—that could serve as potent endogenous clonal markers.

To address these limitations, we present **scLANTERN (single cell Long-reAd liNeage Tracing via Expressed Repeat variatioN)**, a novel, high-throughput methodology for the simultaneous discovery of highly mutable somatic variants and cell-state annotation from full-length single-cell transcriptomes. Utilizing Multiplexed Arrays for Sequencing (MAS-Seq) on the PacBio Revio platform, scLANTERN bypasses the resolution constraints of short-read technologies. The process begins with the efficient isolation of individual cells or nuclei and concludes with the generation of clonal/lineage phylogenetic trees, integrated with detailed annotations of gene isoforms and variants. scLANTERN can be deployed after running canonical short read single-cell profiling with the remaining cDNAs, allowing for data-driven selection of optimal samples. Importantly, this technology can be deployed on primary tissue samples after canonical single cell/nuclei profiling and does not require exogenous genetic barcoding, therefore enabling the establishment of high-resolution lineage tracing of primary tissues during normal development or tumor evolution.

## Introduction

Understanding the lineage relationships among individual cells within a complex organism is fundamental to unraveling developmental processes, tissue regeneration, and the progression of diseases such as cancer (Shapiro et al. 2013). Clonal tracing techniques enable researchers to track the progeny of single cells over time, providing insights into cell fate decisions and lineage hierarchies (K and FM 01/20/2012; McKenna et al. 2016). Traditional clonal tracing methods often rely on genetic labeling strategies using reporter genes or inducible recombination systems (J et al. 11/01/2007; Sanes 1989/01/01; YA et al. 2013 Jul; McKenna et al. 2016). While powerful, these approaches can be limited by the resolution at which distinct clones can be discriminated against and may not capture the full spectrum of cellular diversity. These molecular barcoding perspective lineage tracing methods can only be applied to model cell lines or organisms, not possible for human samples. Recent advances in single-cell sequencing have revolutionized our ability to profile genomic (N et al. 04/07/2011; Tao et al. 2021a; DD et al. 2025 Nov) and transcriptomic variations at the individual cell level (Tang et al. 2009–04–06; MS et al. 2015 Dec). Single cell mitochondria heteroplasmy based lineage tracing has been demonstrated to trace clonal variants in leukemia(LS et al. 03/07/2019). Single cell genomic sequencing has been applied for retrospective lineage tracing (Tao et al. 2021b); however, its application is limited by its throughput and cost, and it remains challenging to integrate both genomic variants and transcriptomics from the same cell. Recently, single cell RNA sequencing has been explored for variant discovery(Zimmermann et al. 2026). However, most single-cell RNA sequencing studies employ short-read sequencing technologies, which may not effectively resolve complex genomic rearrangements or repetitive regions that can serve as unique clonal markers. Long-read sequencing technologies, such as MAS-Seq (Al’Khafaji et al. 2023–06–08) provided by Pacific Biosciences and Oxford Nanopore, produce reads that are significantly longer than those of traditional short-read platforms (Wang et al. 2021 Nov 8; Al’Khafaji et al. 2023–06–08). This allows for more comprehensive detection of structural variants, repetitive elements, and full-length isoforms (Wang et al. 2021 Nov 8; Al’Khafaji et al. 2023–06–08).

In this study, we integrate droplet single-cell isolation techniques with long-read sequencing to perform high-resolution clonal tracing. By analyzing individual cells at the genomic level with long-read sequencing, we can identify unique genetic signatures and reconstruct lineage relationships with unprecedented detail and resolution. Our scLANTERN approach provides a powerful framework for studying clonal dynamics in complex tissues and can be applied to various fields, including developmental biology, stem cell research, and oncology. The ability to trace clones with high resolution opens new avenues for understanding how individual cells contribute to tissue function and disease.

## Materials and Methods

### HAP1 Cell Culture and Ex-Vivo clone preparation

HAP1 MLH1 KO cells (Horizon Discovery, Cat: C699, Lot#37998) were cultured in IMDM supplemented with 10% FBS and 2 mM L-Glutamine at 37°C in 5% CO2. For single-cell deposition, cells were thawed, enumerated, and serially diluted before being plated onto a microfluidic chip (Innovative Biochips LLC) according to the manufacturer’s instructions. Microscopic verification confirmed single-cell occupancy in each well. Subsequently, 200 µL of complete growth medium (IMDM with 10% FBS and 2 mM L-Glutamine) was added to each well. After an initial incubation, the medium was exchanged, and emergent colonies were transferred to a standard cell culture plate for expansion. Discrete colonies were then expanded to establish a second passage, followed by a third passage. Routine maintenance involved weekly complete medium replacements. Finally, third-passage cells were harvested and cryopreserved in a solution of 90% FBS and 10% DMSO for long-term storage in liquid nitrogen vapor.

### HCT 116 Cell Culture and Ex-Vivo clone preparation

The HCT116 cell line (obtained from Genentech gCell, the Genentech internal centralized cell line banking facility), characterized as an adherent, epithelial cell type with an approximate doubling time of 21 hours, was initiated from cryopreserved stocks. Cells were thawed, enumerated, and serially diluted for single-cell deposition onto a microfluidic chip (Innovative Biochips LLC) following the manual. Upon microscopic verification of single-cell occupancy in individual wells, 200 µL of complete growth medium, consisting of RPMI-1640 supplemented with 10% Fetal Bovine Serum (FBS) and 2 mM L-Glutamine, was added to each well. Following an initial incubation period, the medium was exchanged, and emergent colonies were transferred to a standard cell culture plate for expansion. A selection of discrete colonies was subculture to establish a second passage, from which a third passage was subsequently derived. Routine culture maintenance involved a complete medium replacement on a weekly basis. Finally, the third-passage cells were harvested and cryopreserved in a solution composed of 90% FBS and 10% dimethyl sulfoxide (DMSO) for long-term storage in liquid nitrogen vapor.

### Single-Cell Isolation and Ex vivo samples generation with microfluidic device

Single cell clones were generated using a 1CellPlate®-96well microfluidic device (Innovative Biochips LLC). A suspension of viable cells was diluted in complete culture medium to a final concentration of 250–350 cells/mL, and 100 µL of this suspension was loaded into each of the three inlet ports, distributing the cells across the microwell array. Immediately following, the plate was systematically scanned using a bright-field microscope at 10x magnification to identify all microwells containing a single cell. To initiate clonal expansion, 200 µL of pre-warmed culture medium was added to each corresponding outlet well, and the plate was incubated for one to two weeks. Upon confirmation of colony formation, individual clones were harvested and transferred to a standard 96-well plate for a first passage of expansion. After an additional week of growth, the clones were harvested and counted, with the top three lines selected for further propagation based on robust proliferation and desired morphology. These selected clones were then seeded into 6-well plates, cultured to 80–90% confluency, and subsequently harvested and cryopreserved in freezing medium to create master cell banks.

### Barcoding and clonal tracing of LS180 cell line

The LS180 cell line (obtained from Genentech gCell), characterized as a mixed adherent-suspension, epithelial cell type with an approximate doubling time of 27 hours, was initiated from cryopreserved stocks. The TraCe-seq lentivirus barcode library was generated as described in (Chang et al. 2022). For virus transduction, six pools of 500 cells were seeded in a six-well plate and infected overnight with 8 ug/ml polybrene (TR-1003-G, EMD Millipore) at a multiplicity of infection (MOI) of 0.05-0.1. Successfully transduced cells were selected for by culturing in the presence of 2 ug/ml puromycin. Further enrichment of the top 50% of eGFP-expressing cells was performed by FACS on a BD Aria Fusion cell sorter and expanded in cell culture medium supplemented with 2 μg ml–1 puromycin. Barcode diversity was accessed for each of the 6 pools by extracting gDNA for sequencing on the MiSeq. Briefly, TraCe-seq barcodes were amplified using a nested PCR strategy with primers reverse1: AACAGATGGCTGGCAACTAGAAGG*C, forward1: GACGGAAGCGTACAACTGGC*G, reverse2:AATGATACGGCGACCACCGAGATCTACACTCTTTCCCTACACGACGCTCTTCCGATCTACAGT GGATCCACCGAACG*C, foward2: CAAGCAGAAGACGGCATACGAGAT[i7_index]GTGACTGGAGTTCAGACGTGTGCTCTTCCGATCTCGAT GGTCCTGTGCTTCTC*C. The pool with the most barcode diversity and balanced clonal contribution was selected for drug treatment and clonal tracing. LS180 cells transduced with TraCe-seq barcodes were seeded at 8 x 10^5^ in multiple replicates in six-well plates and allowed to attach overnight. The following day, cells were treated with either DMSO or 100 nM Inavolisib, and media containing compounds was replenished every 72h-96h. Cell confluency was tracked throughout the experiment using Incucyte confluence (mm^2^). The DMSO-treated group was collected on day 3 after treatment, whereas the Inavolisib group was collected on day 14 after treatment for scRNA-seq profiling.

### Single cell transcription Library Prep for Hap1 and HCT116 samples

Single-cell transcriptome library preparation was conducted using the 10x Genomics Single Cell 3’ RNA-seq Low Throughput (LT) platform, adhering to standardized manufacturer protocols through the cDNA amplification phase. To enable robust sample multiplexing, we employed hashtag-specific primers during library generation for HCT116. Each clone of HCT116 is counted and pooled equally after hashing. One aliquot of ∼1500 cells total was loaded to one 10x Genomics channel. Another aliquot was further digested with gentle detergent to release nuclei while keeping the hashtag attached according to Dogma-Seq(Mimitou et al. 2021–06–03). The nuclei were counted by Cell Counter and ∼1500 nuclei were loaded to another 10x Genomics channel. Hashtag libraries were synthesized via 12-cycle PCR using 5 µl of pre-amplified cDNA templates, utilizing barcoded primers that are distinct from those designated for the primary cDNA library. Following independent quality control and quantification, cDNA libraries were pooled with their corresponding hashtag libraries at an optimized ratio of 10:1 to ensure balanced read distribution during downstream sequencing.

### PacBio Sequencing

MAS-Seq libraries were prepared in accordance with established protocols. Our study commenced during the pilot phase of MAS-Seq kit development, necessitating the use of two distinct versions. The initial, non-barcoded MAS-Seq Kit was utilized for the preparation of Hap1 ex-vivo samples. These individual libraries were subsequently sequenced using the PacBio Sequel II 8M chip, with one sample allocated per chip, following standard operational procedures. For HCT116 ex-vivo and TraCe-seq samples, we employed the updated MAS-Seq kits, since rebranded as Kinnex. This process began with the hashing of 10x Genomics 3’ cDNA pools. Sequencing was performed using the high-throughput PacBio Revio 25M platform to accommodate the increased scale of these samples.

### Single cell RNA Illumina sequencing data analysis

We use standard cell ranger workflow build following 10xGenomics recommendation to get a cell x gene matrix. We used a customized python script to recover hashtags from hashtag fastqs. We used a customized python script with Salmon to recover TraCe-seq barcodes from fastqs. TraCe-seq genetic barcode recovery from scRNA-seq FASTQ files was performed using Salmon (v. 0.1.1) as described in (Chang et al. 2022).

### MAS-Seq long read data analysis

Raw long-read sequencing data generated via the MAS-Seq workflow were processed using the standardized bioinformatics pipeline recommended by Pacific Biosciences (https://isoseq.how/umi/). Sequence alignment and barcode demultiplexing were performed using default parameters to ensure robust read assignment and data integrity. For cell barcode demultiplexing, we utilized the 10x Genomics 3’ barcode whitelist (3M-february-2018-REVERSE-COMPLEMENTED.txt.gz) and the 5’ barcode whitelist (737K_august_2016.txt.gz). All reads were aligned to the human reference genome (GRCh38), utilizing the Gencode v39 GTF annotation file to ensure precise characterization and annotation of transcriptomic features.

### Variant calling on MAS-Seq long read data

To call short tandem repeats (STRs), we utilized trgt-v2.0.0 (https://github.com/PacificBiosciences/trgt.git) in with the human reference genome hg38 and the “human_GRCh38_no_alt_analysis_set.platinumTRs-v1.0.trgt.bed” comprehensive list for whole-genome STR and repeats, applying all settings as recommended.

To call additional single-nucleotide variants (SNVs) and Indels, we utilized DeepVariant (v1.0.0)(Poplin et al. 2018–09–24), a deep learning-based framework optimized for high-accuracy variant detection. The analysis was aligned to the human reference genome assembly GRCh38 (hg38).

After calling variants, reads were assigned to the allele with the highest alignment score under the Needleman-Wunsch algorithm as implemented in BioPython(Cock et al. 2009 Mar 20). Finally, STRs from TRGT and SNV/Indels from DeepVariant were concatenated into a single count matrix; in case of overlaps between the TRGT and DeepVariant calls, the variant from TRGT was used.

### Pseudo bulk Mutation Calling and Phylogeny Tree Construction

A customized script utilized hashing tags and cell barcodes to group and separate a combined BAM file into individual clone-level (“pseudobulk”) BAMs, with subsequent variant-calling and bulk-level genotyping with TRGT. Subsequently, a Python script merged these clonal VCFs into a character matrix representing the genotypes. This matrix was then filtered to remove non-variable loci and microsatellites whose repeat units were longer than 10 (as these were often miscalled artifacts). Finally, FastTree2 was employed to reconstruct the phylogeny (Price et al. Mar 10, 2010).

### Single cell resolution tree construction

Because single-cell RNA sequencing yields sparse and noisy UMI counts, it is challenging to estimate genotypes at specific loci; therefore, we propose a distance metric based on estimated allelic frequencies within each cell. We define the cellular allele frequency as the fraction of the cell’s transcripts covering the locus that contain an allele – for example, if a diploid cell is heterozygous at a bi-allelic locus and there is no allele-specific expression bias, then its allele frequency at that locus is (0.5, 0.5). Note that in practice, the cellular allele frequency may not be exactly 0, 0.5, or 1, due to allele-specific expression bias, copy number variation, polyploidy, and sequencing errors.

Due to the sparsity of the count data, we smooth the empirical cell allele frequencies by shrinking them towards cluster means, where the clusters are estimated via Expectation-Maximization (EM) of a Multinomial mixture model. This initial clustering is also used to filter for informative loci based on a Mutual Information metric (Rosenberg et al. 2003).

We then construct a phylogenetic distance metric using the squared-Euclidean distance between the cell allele frequencies, or a low-rank approximation thereof computed via PCA. The squared-Euclidean distance is related to the expected edit distance if we sample a random allele from each cell at each locus and is also proportional to Nei’s Minimum Genetic Distance (Nei 1973) and Patterson’s F2 statistic (Patterson et al. 2012) which are commonly used in population genetics to measure population divergences. To induce further smoothing, we use a low-rank approximation of the squared-Euclidean distance by first performing PCA on the cell allele frequencies. Optionally, we include a batch correction method based on projecting out linear discriminants. We then employ the neighbor-joining algorithm (Saitou and Nei 1987) to construct lineage trees from these distances.

We illustrate our approach in **Figure 1c** and provide mathematical details in the Supplementary Note, which includes the EM fitting procedure, the locus selection, batch correction procedure, and theoretical motivation for using the squared-Euclidean distance.

**Figure 1:**
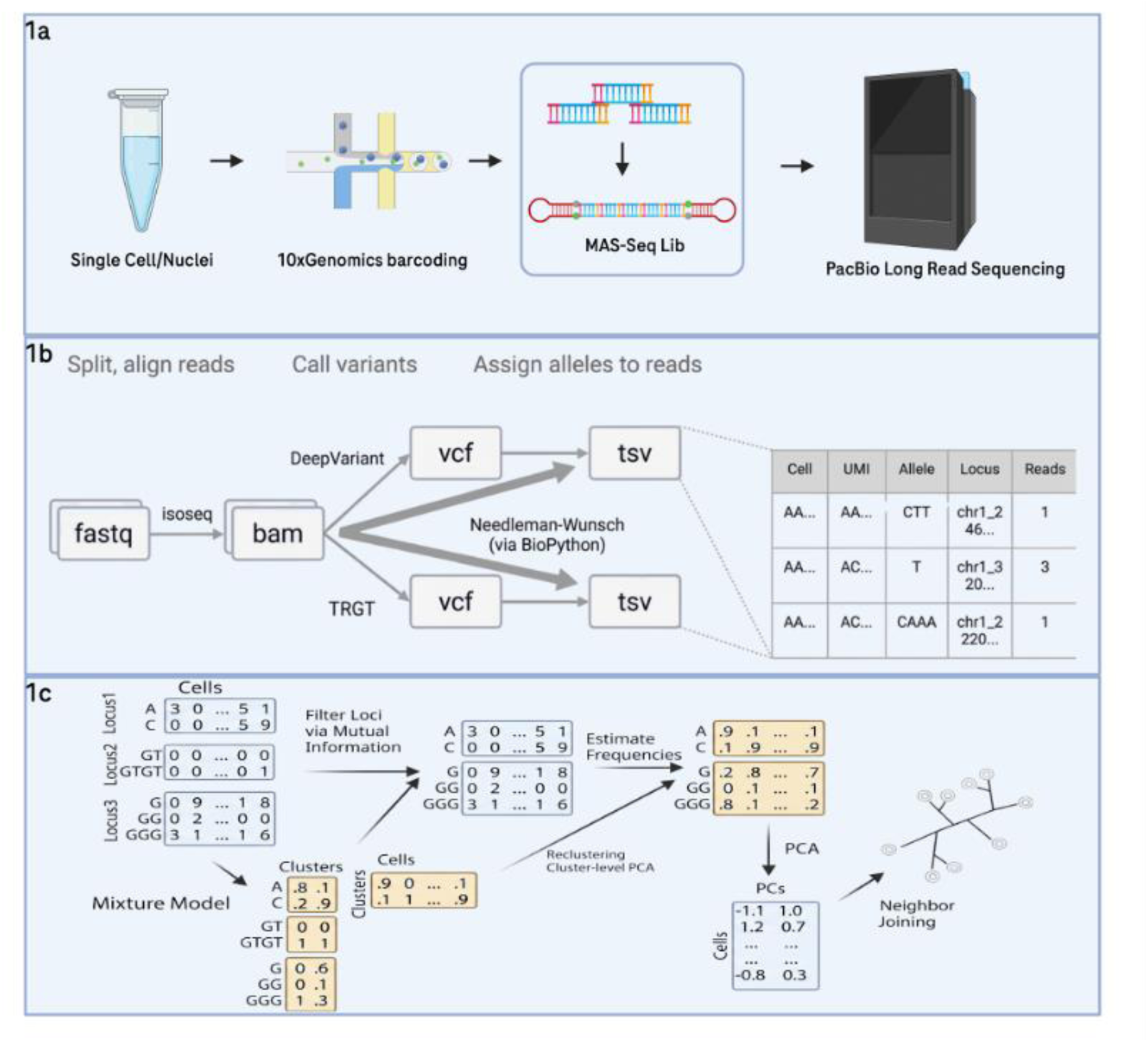
Schematic overview of the single-cell long-read sequencing workflow used for clonal tracing. 1a. Wet lab workflow from cell/nuclei suspension, 10x genomics RNA chip loading, MAS-Seq lib preparation to PacBio HIFI sequencing. 1b. Schematic of bioinformatic pipeline. Reads are split and aligned with PacBio’s isoseq tool to produce BAM files. Then, DeepVariant and TRGT are used to discover variants. Reads from the previous BAMs are then assigned to the best scoring allele by Needleman-Wunsch. Finally, a long data frame for the number of reads per UMI and allele is produced. 1c. Schematic of analysis pipeline. After constructing UMI count matrices per cell, the cells are clustered using a Multinomial mixture model. This initial clustering is used to select informative loci via a rescaled Mutual Information metric. Cell-level allele frequencies are then estimated by shrinking their empirical frequencies towards the cluster means, and a neighbor-joining tree is constructed on the PCA of the allele frequencies.

### Using F_ST_ to identify genes with high mutational burden

For the TraCe-seq data, we used F_ST_, a statistic from population genetics that measures population differentiation (Holsinger and Weir 2009), to identify genes with high mutational divergence between the main expanding and shrinking clones. More specifically, we computed the empirical estimator F_ST_ = (**π**_Between_ - **π**_Within_) / **π**_Between_ for each gene, where **π**_Within,_ **π**_Between_ are the average number of pairwise differences for reads within and between populations (Hudson et al 1992). The pairwise differences were in turn computed as the number of variant loci minus the inner product of the allele frequencies, i.e. **π**_Between_ = *L* − *P*_1_ ⋅ *P*_2_ and 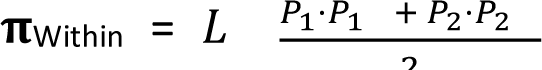 where *P*, *P* are the allele frequencies of clones 1 and 2. *P* and *P* were in turn estimated using the empirical allele frequencies, shrunken with a pseudocount of 5 towards prior frequencies from the Multinomial mixture model fit by scLANTERN (where the prior frequencies are obtained by averaging the mixture model’s expected frequencies across all the cells within each subpopulation).

### Using Adjusted Rand Index (ARI) to quantify PCA-based phylogenetic distance metric

To quantify the agreement of our PCA-based phylogenetic distance metric with the ground truth clones, we obtained clusters using Ward’s minimum variance criterion and computed the Adjusted Rand Index (ARI), using the same number of clusters as clones (5 in the HAP1 dataset, 6 in the HCT116 dataset).

## Results

### scLANTERN experimental and computational workflow

We developed scLANTERN as a full solution, which includes both experimental protocols and the supporting computational pipeline. The experimental workflow is shown in **Figure 1a** as a comprehensive methodology for generating high-resolution single cell data and starts with the isolation of single cells or nuclei from diverse biological materials such as fresh or frozen tissues and cell suspensions. Following optional hashing for sample multiplexing, cells/nuclei are encapsulated using 10x Genomics technology, where reverse transcription converts mRNA to full-length cDNA. This cDNA undergoes pre-amplification, with optional hash primer spike-in, followed by purification and quantification. Subsequently, 15 ng of purified cDNA is utilized for MAS-Seq library construction, involving further amplification, dimer removal, 16 PCR cycles with specialized primers, digestion, ligation into 22kb fragments, and circularization with PacBio dumbbell adaptors. The resulting circular library is then subjected to PacBio sequencing, yielding long reads that are processed through a dedicated single-cell MAS-Seq pipeline. This pipeline breaks down long reads into individual full-length cDNAs, extracts UMIs and cell barcodes, aligns sequences to a reference genome for annotation, and enables diverse downstream analyses including gene expression quantification, cell type identification, mutation discovery, and lineage inference, thus providing a powerful tool for unraveling cellular heterogeneity and complex biological processes at single-cell resolution.

The bioinformatic process for mutation detection and subsequent inference of cellular lineages is implemented in a Nextflow pipeline (“sclantern-nf”), which is illustrated in **Figure 1b**. The initial step is the alignment of full-length cDNA BAM files, where each individual read is initially enriched with a unique molecular identifier (UMI) and cell barcodes, extracted directly from the BAM tags. Following this initial tagging, the reads undergo re-alignment to the reference genome using Minimap2. Post-alignment, the Genome Analysis Toolkit (GATK) is leveraged to split the reads into their respective exonic and intronic regions.

The tagged and exon-intron split BAM file then serves as input for variant calling algorithms. DeepVariant (Poplin et al. 2018–09–24), a neural network-based variant caller, is applied for the identification of single nucleotide variants (SNVs) and indels. Concurrently, TRGT (Tool for Rapid Genotyping of Tandem Repeats) is utilized for genotyping short tandem repeats (STRs), which are often challenging to call with standard variant callers. After the initial variant calling with DeepVariant and TRGT, each read at each variant is assigned to the allele with the highest alignment score under the Needleman-Wunsch algorithm as implemented in BioPython(Cock et al. 2009 Mar 20).

Once variants alleles are assigned to every UMI, an initial clustering of the cells is generated via a multinomial mixture model, which is used to smooth cell-level allele frequencies for PCA and Neighbor-Joining, as described in Materials & Methods (“Single cell resolution tree construction”). This pipeline is shown in **Figure 1c** and implemented in our sclanternR R package, which also includes plotting functionality to visualize the resulting lineage inferences.

### scLANTERN successfully deconvolves single cell clonal evolution of the haploid HAP1 chronic myeloid leukemia cell line

To validate the precision and reliability of scLANTERN to infer clonal evolution, we initially selected the Hap1 cell line, since this model is mostly haploid, with variants typically originating from only one chromosome. As illustrated in **Figure 2a**, *in vitro* Hap1 single-cell clones were generated with known progeny relationships by growing them from single cell to single cell clone, picking a single cell from the parent clone, and culturing it to generate offspring of single cell clones. Each clone was named by their generation and culture position in the plates. We then loaded ∼500 cells per clone to 10x Genomics Low Throughput 3’ RNA workflow to generate sufficient coverage per cell and sequenced the full-length cDNA using the MAS-Seq protocol with PacBio HiFi reads. Subsequent application of our mutation discovery pipeline led to the identification of thousands of Short Tandem Repeats (STRs) and Single Nucleotide Variants (SNVs). A maximum likelihood tree at the clonal pseudo bulk level agreed with the ground truth tree (**Figure 2b**). At the single-cell level, a PCA of the estimated allele frequencies of the top 50 informative loci largely recapitulated the ground truth phylogeny, with cells from clones D7A, D7B, and E4B1 at the tips of the tree, and cells from the parent clones D and E in the internal branches (Figure 2e); the neighbor-joining tree also recapitulated this structure (Figure 2f), and the ground truth clones had an Adjusted Rand Index (ARI) of 0.42 when compared with clusters based on Ward’s minimum variance criteria in the PCA space.

**Figure 2:**
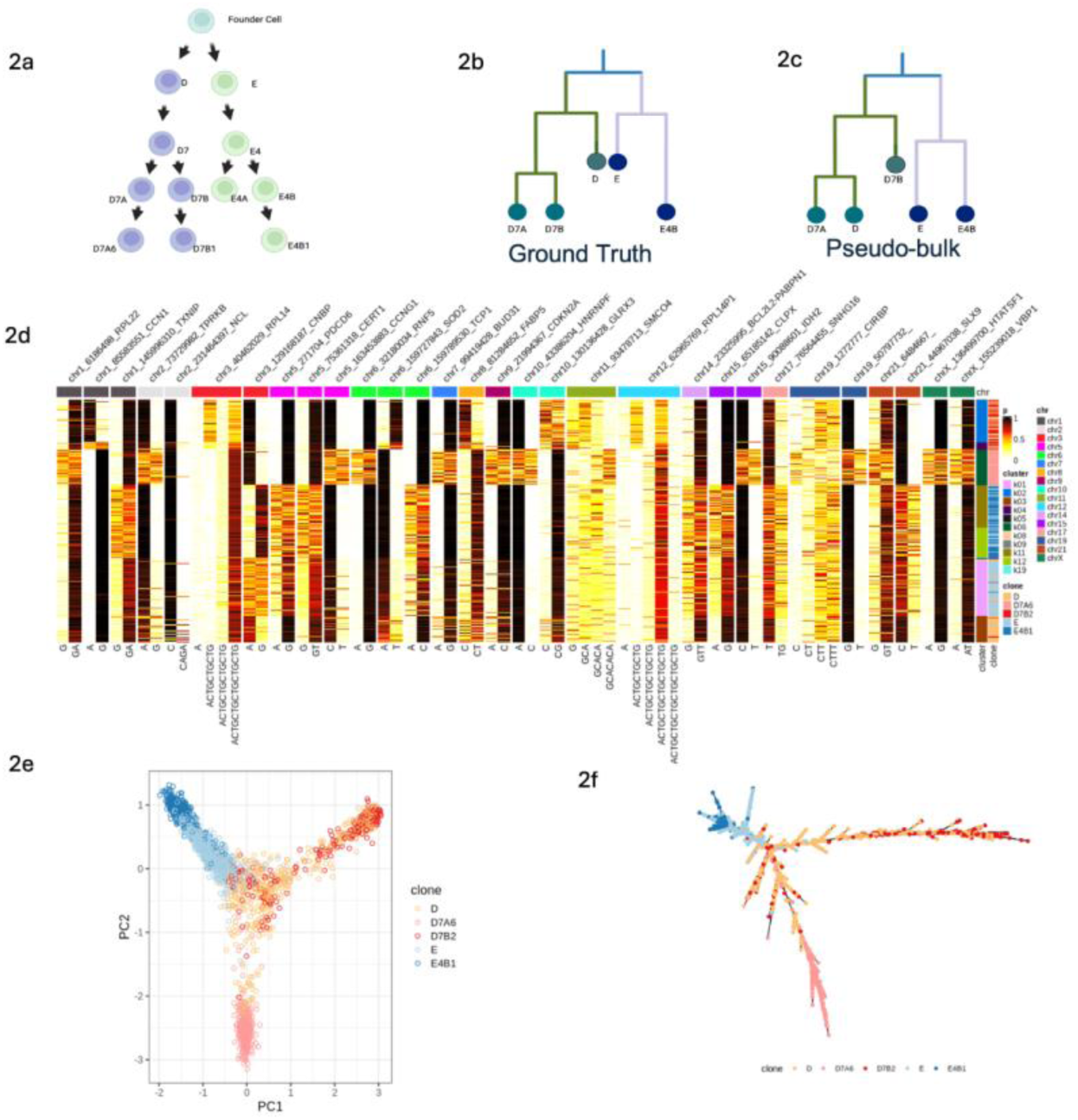
Mutation based Phylogenetic Tree reveals progeny relations in Haploid ex vivo clones and single cells. 2a. Single cell clones are prepared from a single cell, and subcloning, culturing and banking document their generations. 2b. Ground truth phylogeny tree for selected clones. 2c. a phylogenetic tree reconstructed based on pseudobulk clonal variants from TRGT. 2d. Heatmap of estimated allele frequencies at the top informative loci. Cells are annotated by the ground truth clone as well as the cluster assigned from the multinomial mixture model. Loci were selected using a mutual information metric of the allele frequencies with the mixture model clusters. 2e. PCA of estimated allele frequencies at the top 50 loci. 2f. Single cell neighbor-joining tree based on the square-Euclidean distance of the PCA loadings.

f

### scLANTERN confirms the clonal and single cell phylogenetic evolution of the HCT116 colorectal cancer cell line

To assess the performance of our scLANTERN approach in a more clinically relevant model, we utilized the diploid colorectal cancer cell line HCT116. Single-cell clones were generated and banked using an approach analogous to that employed for the Hap1 line, with lineage relationships documented to establish experimental ground truth (**Figure 3a**). Six clonal lineages were selected for analysis, and a ground-truth phylogenetic tree was constructed based on culture-derived lineage relationships (**Figure 3b**). Since STRs are also present in intronic gene regions, we also generated matched single nuclei profiling, since this strategy would enable using frozen samples and enriching for introns, therefore potentially increasing the applicability and resolution of our technology. To this end, we initially hashed and pooled equal counts of cells from each clone and took an aliquot to load ∼1500 cells for 10x Genomics Low Throughput 3’ RNA workflow. Then another aliquot of the same pool was digested with gentle detergent to release the nuclei (following the Dogma-Seq protocol), and ∼1500 nuclei were loaded to another 10x Genomics 3’RNA channel. After sequencing, scisoseq recovered 2x more nuclei than whole cells (1269 vs 676), with the nuclei libraries having a lower percentage of reads in cells compared to whole cell libraries (69% vs 91%), resulting in 3x more UMIs per whole cell vs single nuclei (median 32,000 vs 98,000 UMIs per cell, **Sup Fig S5**).

**Figure 3:**
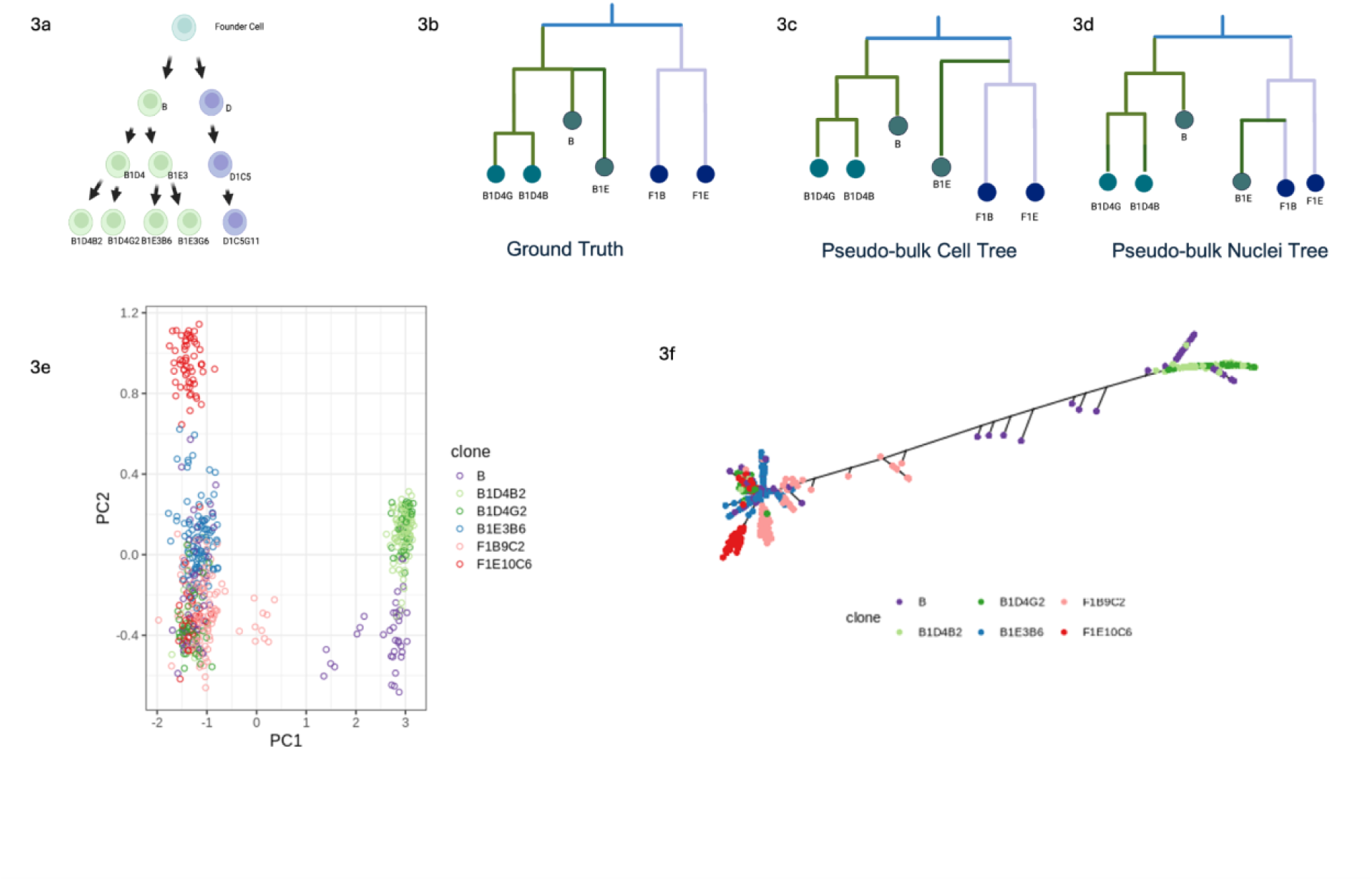
Reconstructed lineage relationships among single cells based on mutations for HCT116 cancer cells. 3a. Single cell clones’ generation with known ground truth progeny relations. 3b. Ground truth of selected clones. 3c. Pseudo-bulk nuclei phylogeny tree reconstruction with STR variants 3d. Pseudo-bulk cell phylogeny tree reconstruction with STR variants. 3e. PCA of estimated allele frequencies (cells only) 3f. Single cell clonal lineage tree reconstruction by neighbor joining (cells only)

Pseudobulking cells by library prep (whole-cell or nuclei) and ground truth clone yielded maximum-likelihood trees that placed whole-cell and nuclei pseudobulks from the same clone together (**Sup Fig S4**), while recapitulating much of the topological structure of the true phylogeny, such as the grouping of clones B1D4B2 and B1D4G2. The pseudobulk tree built from the cell library prep was similar to the pseudobulk tree built from the nuclei library prep (**Figure 3b, 3c, 3d**). At the level of individual cells/nuclei, we computed phylogenetic distances from a PCA of the estimated allele frequencies at informative loci, performing separate cell-only, nuclei-only, and combined analyses; for the combined analysis, we included batch correction between the cell/nuclei libraries. The cell-only analysis had a good concordance of the PCA embedding with the ground truth clones (**Figure 3e, 3f**), with Ward minimum-variance clusters having an ARI of 0.38. The nuclei-only and combined analyses successfully clustered clones B1D4B2 and B1D4G2 together (**Sup Figure S3**) but otherwise did not perform as well (ARI of 0.1 and .09 respectively), possibly due to lower sequencing depth of the nuclei. In both single-cell and pseudobulk analyses, clones B, B1D4B2, B1D4G2 tended to group together, while F1B9C2, F1E10C6, and B1EB6 formed another group; the misclustering of B1EB6 highlights the challenges of mutation calling in this sparse context, suggesting a need for further development (such as with improved variant calling algorithms or by combining the RNAseq data with other modalities).

### scLANTERN successfully reconstructs longitudinal clonal pairs in LS180 cells with built-in TraCe-seq genetic barcodes and identifies expanding and shrinking clones

To further evaluate the performance of the scLANTERN pipeline, we utilized TraCe-seq, a methodology employing genome-edited genetic barcodes to provide ground-truth lineage labels(Chang et al. 2022). We applied this approach to the highly mutable colorectal cancer cell line LS180, which, while polyclonal in nature, exhibits bottleneck dynamics under therapeutic selective pressure. Since LS180 cells harbor a mutation in the PI3Ka gene, we treated this model with 100nM concentration of Inavolisib (Hanan et al. December 1, 2022), a selective PI3Ka inhibitor, for 14 days and performed longitudinal sampling of the tumor cells at baseline (Day 0) and Day 14 for both TraCe-seq and scLANTERN lineage profiling. In addition to validating our lineage tracing technologies, this approach may point to molecular mechanisms underlying the emergence of Inavolisib resistant clones at day 14.

Pseudo-bulk phylogenetic analysis of the long-read sequencing data demonstrated robust concordance between mutation-based lineage reconstructions and the established TraCe-seq barcode assignments; with 13 out of 15 clone pairs harboring the same eGFP barcode between baseline and day 14 clustering together in the phylogeny reconstructed by FastTree2 on expressed STRs (**Figure 4a**). At the single cell level, this dataset was more challenging for clonal reconstruction compared to our ex vivo HAP1 and HCT116 clonal models, due to the higher number of cells (∼10,000) resulting in lower sequencing depth and greater sparsity per cell. Nevertheless, the PCA of the estimated allele frequencies (with batch correction for timepoint) largely clustered cells according to their clonal origins (**Figure 4b**, **Supplemental Fig S2**). A neighbor-joining tree of cells from the 8 biggest clones largely recovered clusters for 2 clones and partially recovered a third clone as a single cluster (**Figure 4c**). Finally, differential analysis of STR mutation enrichment between the longitudinal time points identified specific mutation burdens localized within pathways targeted by the therapeutic agent (**Figure 4e**). The two clones that scLANTERN performed best on were clone group 95187 and clone group 90794. Clone 95187 expanded across timepoints, while clone 90794 shrunk (**Figure 4d**), but note that these clones each had cells in both timepoints, which scLANTERN clustered by clone instead of by timepoint (**Supplemental Figure S2**). We further compared the mutations discovered from these two clones between day 0 baseline and day 14 under 100nM Inavolisib treatment and found significant enrichment at day 14 cells for pre-existing variants of genes known to modulate the PI3K pathway, which warrant future functional validation (**Figure 4e**).

**Figure 4:**
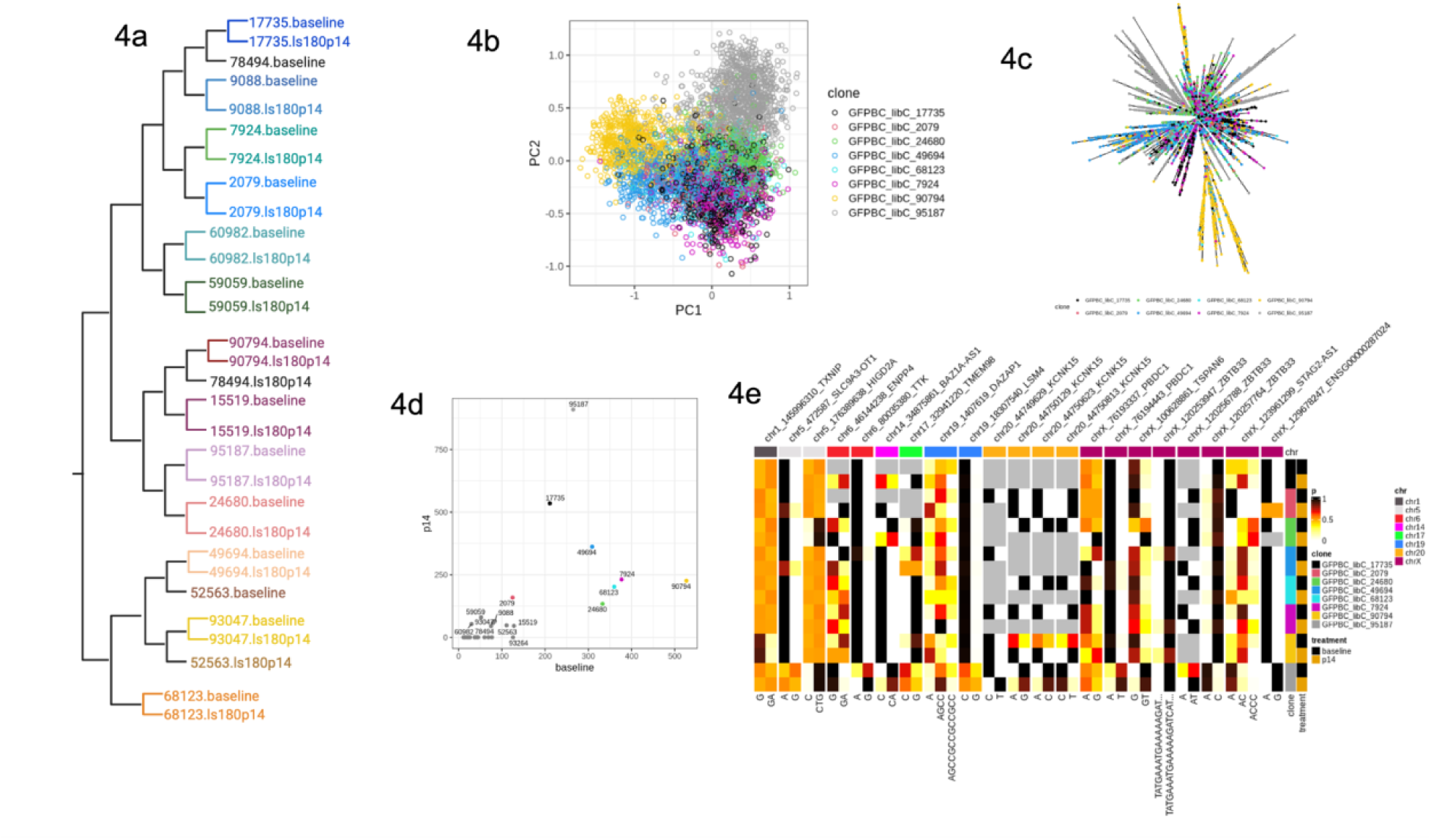
Reconstructed lineage relationships among single cells based on mutations for with built in clonal tags. 4a. Pseudobulk clones pairing between baseline and day14 based on STR variants. 4b. PCA of estimated allele frequencies. 4c. Neighbor joining tree, subsampling 100 cells per (clone, day) from the top 8 clones. 4d. Clonal abundances (number of cells) at each of the two timepoints. The top 8 abundant clones are colored. Clone 95187 has the greatest expansion, while clone 90794 has the greatest reduction between the timepoints. 4e. Pseudobulk (clone+day) allele frequencies. We show the top 15 genes with greatest divergence (F_ST_) between the expanding and shrinking clones (95187, 90794), using the smaller of the two F_ST_ values for baseline, day14 when sorting. Gray tiles indicate NA (0/0), which occur when a pseudobulk sample has no reads at the locus.

Altogether, these findings demonstrate the utility of scLANTERN in resolving clonal selection without barcodes to investigate mechanisms of therapeutic adaptation, establishing a framework for future studies employing deeper sequencing to optimize single-cell resolution lineage tracing.

## Discussion

In this work we validated our scLANTERN lineage tracing technology using the HAP1 and HCT116 cell line models with known ground truth of phylogeny relationship among single cell clones, demonstrating its capacity to reconstruct phylogenetic trees at both pseudo-bulk and individual cell resolutions as proof of concepts. In HAP1 clones, scLANTERN identified thousands of STRs and SNVs, which are further ranked to estimate the most informative top variants for tree construction. The resulting phylogenetic reconstructions demonstrated very good concordance with known ground-truth clonal relationships. For the HCT116 colorectal cancer cell line, scLANTERN was able to cluster cells according to their clonal origins, though it performed better on whole cells than nuclei, possibly due to higher coverage (**Sup Fig S5**). Extending our application to TraCe-seq samples in the LS180 cell line, scLANTERN was able to recover much of the TraCe-seq clonal structure across two timepoints, especially in the most prominent expanding or shrinking clones, and provide clone-specific enrichment of gene variants that could play functional roles in the emergence of these resistant clones.

Importantly, scLANTERN is compatible with intact nuclei or fresh cells from patient tumor samples and can be readily applied to already existing full-length single-cell cDNA libraries generated via 10x Genomics pipelines. This allows a “plug and play” workflow for critical tissue samples, with an initial profiling by canonical single cell/nuclei RNA or multiome technologies to establish biological hypotheses, and the subsequent deployment of scLANTERN on the same cDNA for lineage reconstruction. In addition, and unlike traditional methods which rely on single-cell DNA sequencing and are often limited by high costs and low throughput (N et al. 04/07/2011; CJ and K 2020 Apr; Tao et al. 2021a; Neumeier et al. 2025), our approach leverages the information-rich nature of full-length transcripts—including gene isoforms and variants—to provide a comprehensive understanding of cell and clone evolutionary stages, and their functional association with specific gene alterations.

While our study demonstrates the utility of scLANTERN, it is also subject to some limitations. The discriminatory power of this method depends on the availability and sequencing coverage of *de novo* somatic mutations; consequently, it may yield insufficient resolution for cells from tissues with low mutational burdens, especially in sparsely sequenced samples. This challenge was evident in our LS180 TraCe-seq dataset, where we pooled ∼10K TraCe-seq cells in one PacBio Revio run, in comparison to ∼2K HCT116 single cell or nuclei pool in three PacBio Revio runs. While we successfully reconstructed clonal pairings across time points, the single-cell resolution trees exhibited noise attributable to this sparse coverage. To mitigate these limitations, future implementations could incorporate targeted enrichment panels to reduce the cost per mutation and increase the number of cells analyzed per sequencing run. Alternatively, the depletion of highly expressed housekeeping genes—for instance, via *in vitro* CRISPR-based approaches— could increase the depth of coverage for lower-abundance transcripts.

The integration of high-resolution mutation detection and deep transcriptomic characterization within a single assay allows for the direct linkage of specific genomic alterations to distinct cellular phenotypes. Our findings establish that long-read RNA sequencing provides a robust and scalable framework for investigating clonal dynamics within complex biological systems and clinical tissues. By unlocking the informational potential of repetitive genomic regions, scLANTERN offers a powerful new avenue for research in oncology, immunology, and developmental biology. However, computational challenges regarding the assignment of mutations to single cells in ultra-sparse data remain. Although recent bioRxiv studies (Lougheed et al. 2025) have investigated STR mutation discovery in long-read RNA sequencing, and our work extends these efforts to single-cell resolution, further studies are needed to improve the algorithm handling sparse single cell data. We believe that the continued evolution of long-read sequencing throughput, such as reusable nano pore chips (Kokoris et al. 2025–03–15) (Roche SBX, PacBio) and improved chemistry (PacBio 2026), combined with the development of targeted enrichment methods for the most variable STR, hold the promise to further scale this technology for routine clinical diagnostics and precision medicine.

In conclusion, our work offers a proof-of-concept demonstration that expressed repetitive regions represent a valuable resource for mining somatic variants and facilitating lineage tracing. Its pairing with the identification of gene alterations and variants, provides unprecedented potential for understanding their impact on cellular function in multiple developmental and disease contexts.

## Authors’ Contributions

N.R supervised the study; L.T designed and executed the experiment; J.K led the statistical model development; YT. F developed the bioinformatic pipeline and an early version of the statistical model; D.N provided the TraCe-seq libs and analysis supporting; J.K and L.T led the analysis, L.T, J.K, D.N, N.R wrote the manuscript, L.T and N.R oversaw the resources.

## Acknowledgments

The authors express their gratitude to the Genentech FACS core for their essential role in cell sorting and to Genentech gCell for their assistance with cell line banking. We also acknowledge the contributions of Roche Diagnosis and MedGenome for their PacBio sequencing services, which were vital to this research.

## Data Availability

Upon publication, all data will be made accessible through the NCBI public repository.

## Code Availability

The code used in this study will be made available at:

● https://github.com/genentech/sclanternR
● https://github.com/genentech/sclantern-nf

## Ethics declarations

The human cell lines utilized in this study were acquired from both commercial and centralized repository sources. The HAP1 cell line, carrying a targeted knockout of the DNA mismatch repair (MLH1) gene, was purchased commercially from Horizon Discovery (Waterbeach, UK). The human colorectal carcinoma cell lines HCT116 and LS180 were obtained via gCELL, the centralized cell repository at Genentech, Inc. (South San Francisco, CA, USA)

---

**Sup Figure 1:**
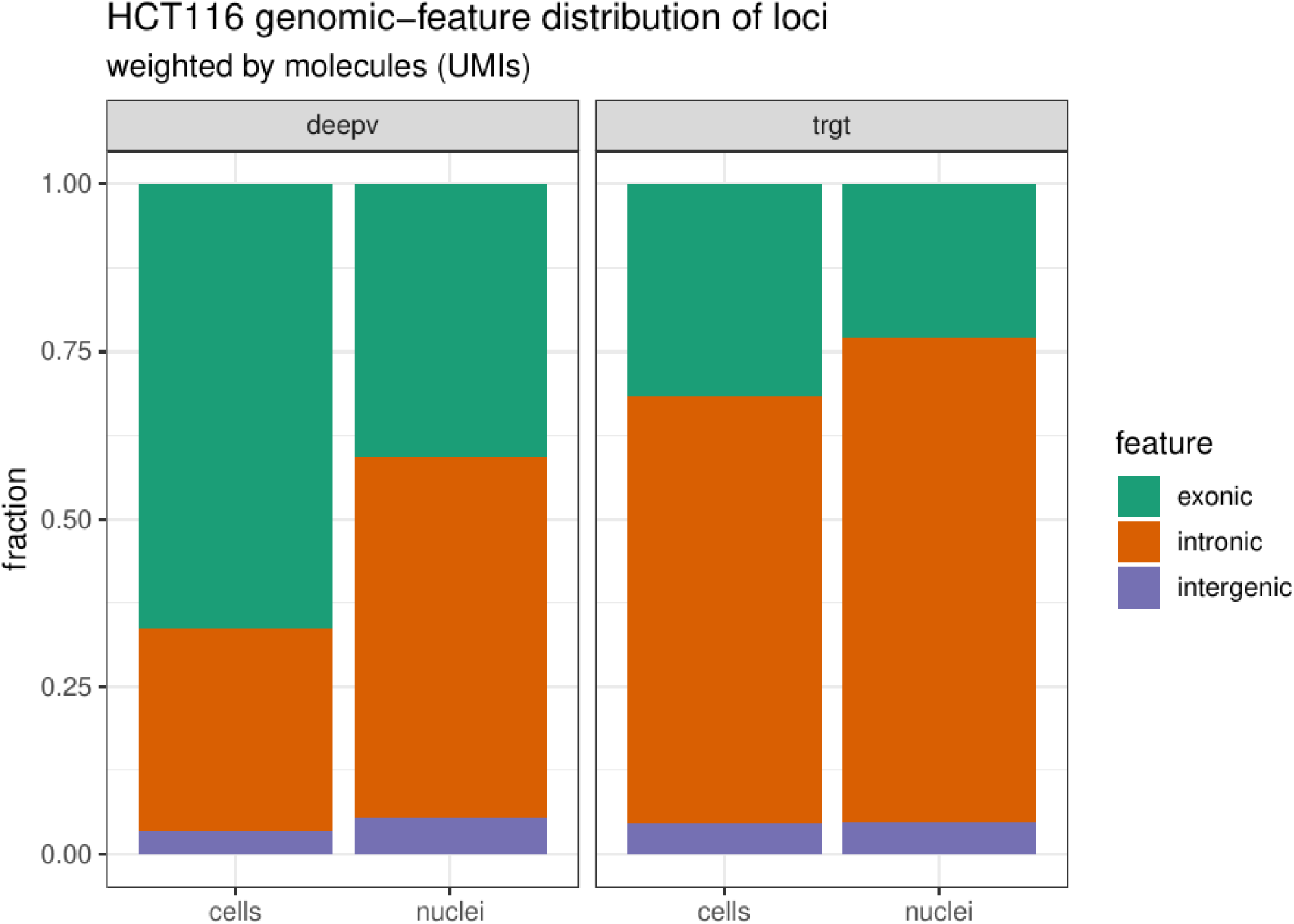
Loci distribution between HCT116 cell dataset and nuclei datasets. Sup Fig1: HCT116 variations distribution between cells and nuclei

**Sup Figure 2:**
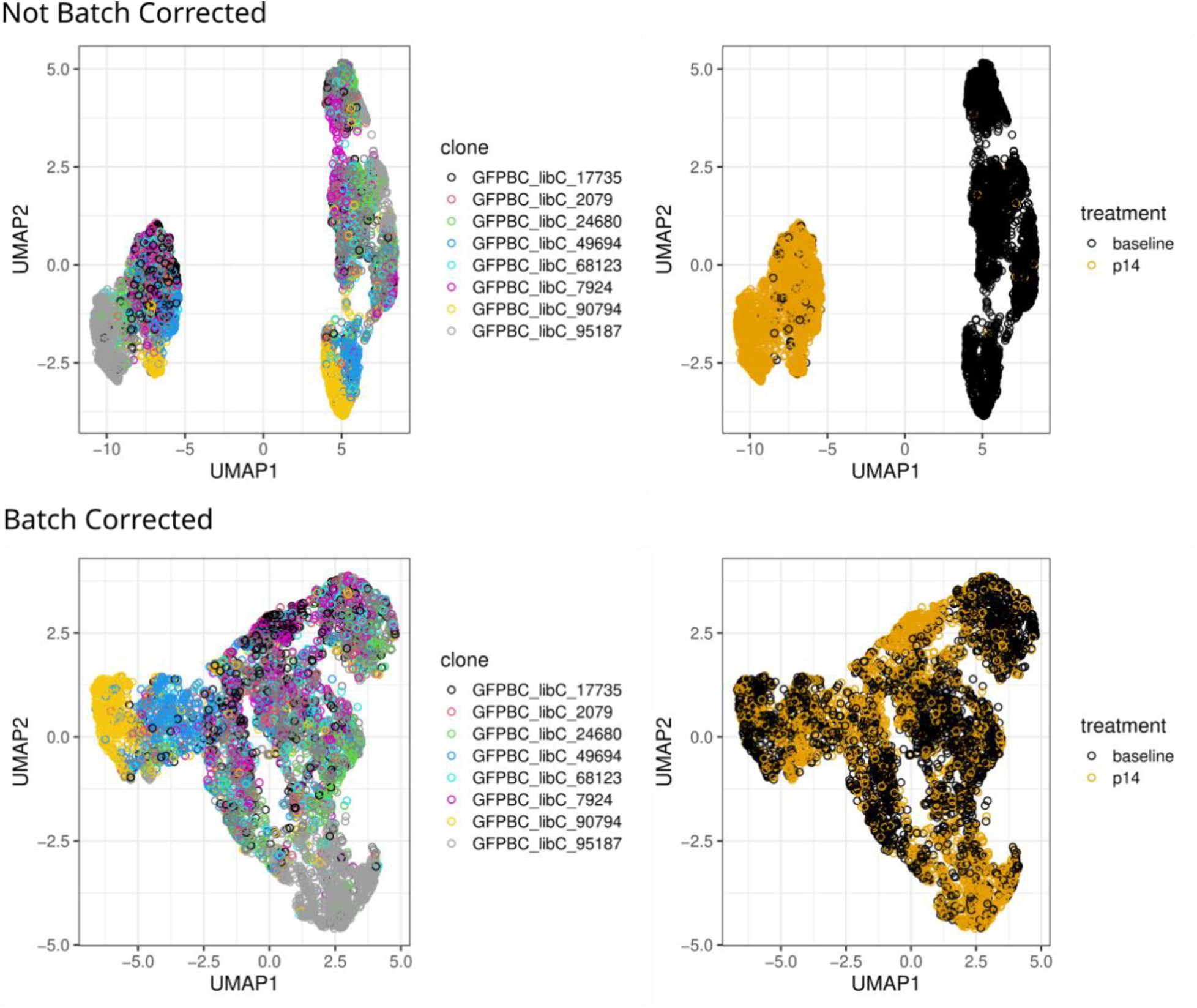
UMAP of PCA loadings of TraCe-seq dataset, with and without batch correction. Sup Fig 2: UMAP of PCA loadings of TraCe-seq dataset, with and without batch correction. Applying the batch correction for the condition (baseline vs day14) prevents the cells from clustering by condition, and instead they cluster by clone. The batch correction works by projecting out the linear discriminant for the condition in the PCA space.

**Sup Figure 3:**
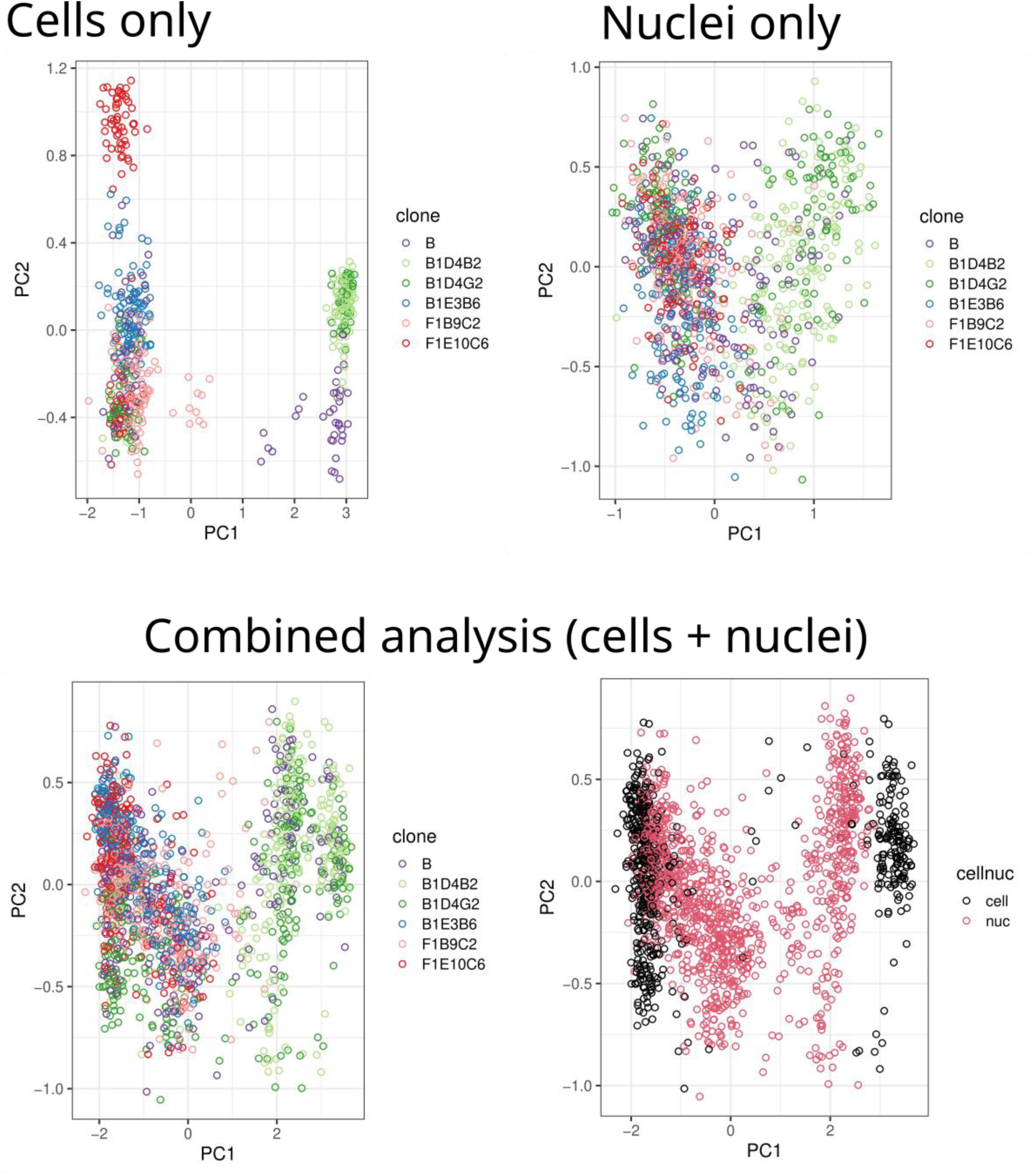
HCT116 cell-only, nuclei-only, combined analysis. Sup Fig 3: PCA from analyses on whole cells only, nuclei only, and combined (cells + nuclei). The whole-cell analysis had the best separation of clonal groups; the nuclei and combined analyses separated the B1D clones but failed to recover finer structure. Despite performing the linear-discriminant based batch correction, the combined analysis still retained some separation between the whole-cell and nuclei clusters.

**Sup Figure 4:**
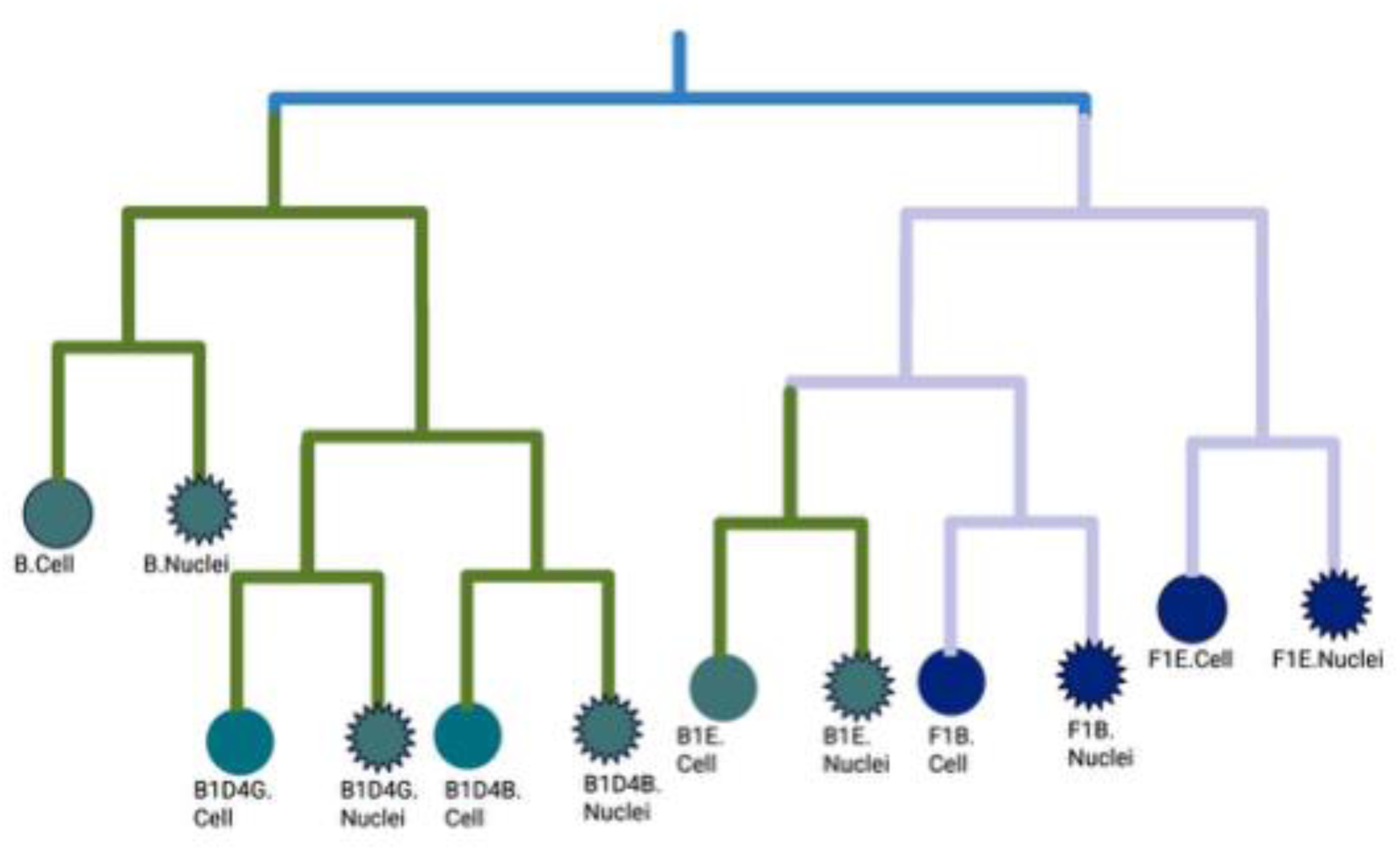
HCT116 Cell lib pre and Nuclei lib pre joint pseudobulk tree. Sup Fig 4: Same clones from HCT116 Cell lib pre and Nuclei lib pre are grouped togethe

